# Evaluating pesticide mixtures for resistance management in asexual insect pests

**DOI:** 10.64898/2026.08.24.746641

**Authors:** Luna Qingyang Li, Ricardo Kanitz, Philip Madgwick

## Abstract

Many economically important insect pests reproduce through asexual or partially asexual life cycles, yet how reproductive mode influences insecticide resistance management remains unclear. The choice of resistance management strategy has been suggested to differ for sexual and asexual pests. For instance, current IRAC guidance suggests that pesticide mixtures are less effective in non-mating pests than in sexually reproducing populations. Here, stochastic evolutionary simulations are used to compare resistance evolution under sequences and mixtures across four reproductive modes observed in pests of economic importance: sexual reproduction, obligate parthenogenesis, cyclical parthenogenesis and haplodiploidy. Contrary to current expectations, mixtures are not disadvantaged in asexual populations and, in some cases, lead to delayed resistance evolution compared to sexually reproducing populations. These differences arise as the result of reduced genetic recombination which constrained the assembly and spread of multi-resistant genotypes. Overall, these findings suggest that mixtures remain a viable resistance management strategy for pests with asexual reproduction.

## Introduction

Although the majority of insect pests of economic importance reproduce sexually, some of the most problematic pests for resistance management exhibit traits of asexuality (Adesanya et al., 2021). Among insect pests, two forms of asexuality dominates: first, cyclical parthenogenesis, found in multiple aphid species, where long periods of parthenogenesis are punctuated by occasional sexual mating on a seasonal basis (Simon et al., 2002); and second, haplodiploidy, common in species of thrips (Krueger et al., 2017), mites (Tuan et al., 2016) and whiteflies (Byrne & Devonshire, 1996), where unfertilised eggs give rise to haploid males and fertilised eggs become diploid females. Due to their generalist nature and high reproductive capacity, pests capable of asexual reproduction cause significant damage to economically valuable crops (Simmons et al., 2008; Mathers et al., 2017; Adesanya et al., 2021; Singh et al., 2021; Perier et al., 2022). Control of these pest species is dominated by the use of synthetic pesticides (Mouden et al., 2017; Horowitz et al., 2020; Singh et al., 2021). Across species, it is generally recommended to rotate multiple chemicals with distinct modes of action as a resistance management strategy (Wu et al., 2019; Horowitz et al., 2020; Perier et al., 2022). If possible, chemicals should be selected according to the resistance status of the local population (Adesanya et al., 2021). However, extensive pesticide resistance in certain species, such as the two-spotted spider mite (*Tetranychus urticae*) and the peach potato aphid (*Myzus persicae*), both of which display resistance to most classes of agrochemicals currently in use (Adesanya et al., 2021; Singh et al., 2021), leaves few compounds efficacious for pest control.

The recommended strategies for insecticide resistance management in these asexual pests are based on limited evidence from resistance modelling, with little emphasis on the influence of reproductive mode on resistance evolution. A majority of theoretical and computational work up to date has focused on sexually reproducing populations (Comins, 1986; Roush, 1989; Barbosa, 2012; Levick et al., 2017; South & Hastings, 2018; Madgwick & Kanitz, 2022b; Hobbs et al., 2023; Madgwick et al., 2024). Using analytical techniques, Mani showed in 1985 that increasing the rate of recombination from 0 to 0.5 retarded the spread of resistance (Mani, 1985), suggesting that beneficial alleles can sweep through clonally reproducing populations more rapidly than in sexual populations. A more recent computational analysis, based on selection phase dynamics, showed that mixtures are less preferable to sequences and rotations in asexual pests (Helps et al., 2020). In line with such evidence, the current IRAC guidance notes that mixtures provide less benefit on managing resistance in non-mating pests (IRAC, 2023).

In evolutionary terms, prior evidence has shown that asexual reproductive systems can profoundly influence resistance evolution. While parthenogenesis may accelerate the spread and selection of resistance alleles (Carrière, 2003; Rubio-Melendez et al., 2019; Bendall et al., 2022; Pakrashi et al., 2025), the lack of recombination can constrain long-term adaptation through clonal interference (Gerrish & Lenski, 1998; Fujita et al., 2020). Evidence from other clonal systems supports the concurrent application of multiple compounds as a successful resistance management strategy. For example, combination therapies are well-established for their ability to control clonally replicating populations ranging from bacteria (Woods & Read, 2023) to viruses (Gibas et al., 2022) to cancer cells (Jin et al., 2023). Although each of these clonal systems has unique characteristics, such as the presence of horizontal gene transfer in bacteria (Arnold et al., 2022) and intratumor heterogeneity in cancers (Dagogo-Jack & Shaw, 2018; Marusyk et al., 2020), there is strong empirical evidence that compound combinations can be effective at controlling clonal populations. These observations stands in contrast with theoretical literature in pest management, where mixtures are currently not a recommended strategy. One plausible explanation for this discrepancy lies in the in the exclusion of emergence phase dynamics in current pesticide resistance modelling studies (Mani, 1985; Helps et al., 2020). In clonal systems such as bacteria and viruses, combination therapies are deployed for their ability to suppress the emergence rather than the selection of multi-resistance genotypes (MacLean et al., 2010). Given the ease at which beneficial genotypes can rapidly sweep through clonal populations, it is likely that managing resistance in clonal populations depends critically on lengthening the emergence phase. Even in sexually reproducing pest populations, theoretical work has demonstrated the importance of understanding emergence phase dynamics in achieving long-term pest control (Madgwick et al., 2024), yet there has been no equivalent analysis for asexually reproducing pests.

Although most asexual insect pest populations are not exclusively clonal, insights from managing clonal populations in other contexts is currently at odds with recommendations for agricultural practices. Resolving this discrepancy requires a better understanding of resistance evolution in asexually reproducing pests. Here, extensive computational simulations are used to examine the emergence and selection phase dynamics of monogenic resistance under three reproductive modes – obligate parthenogenesis or asexual reproduction, cyclical parthenogenesis, and haplodiploidy – and compare them with an otherwise equivalent sexually reproducing population. The simulations assess the effectiveness of sequences and mixtures for delaying resistance evolution and identify reproductive mode-specific dynamics relevant to resistance management. Together, these findings will help inform resistance management strategies for asexually reproducing pests.

## Methods

Wright-Fisher simulations (Fisher, 1930; Wright, 1931) are used to investigate how reproductive strategy influences resistance evolution under single- and dual-compound selection (sequences vs. mixtures). Separate population genetic models are implemented for sexual reproduction, asexual reproduction, cyclical parthenogenesis, and haplodiploidy. Resistance to each compound is assumed to arise from a *de novo* monogenic allele. Genotype fitness values under selection follows prior convention (Levick et al., 2017; Madgwick & Kanitz, 2022b). In haplodiploidy, haploid males have the same fitness as the corresponding homozygous female.

A total of 10,000 randomly sampled parameter sets with beneficial resistance alleles are chosen for stochastic simulations. For each reproductive mode, sequence and mixture strategies are simulated with failure thresholds of either 25% or 75%, producing four datasets. For each parameter set within each condition, 10,000 simulations are run for up to 2,000 generations or until both resistance alleles reached the failure threshold. For each allele, the simulation records the number of generations taken until heterozygote emergence, homozygote emergence, and compound failure. Simulation outcomes are compared using difference distributions between conditions. All simulations and analyses are performed in R.

### Parameter sampling

Simulation parameters apart from mutation rate are sampled according to Table 1. All parameters with 0 – 1 range are randomly sampled from a uniform distribution. Otherwise, the exponent is randomly sampled from a uniform distribution. Mutation rate (*m*) is sampled relative to the population size. The number of *de novo* mutations per diploid genome per generation (*N_m_*) is sampled from a uniform distribution between 0.05 and 0.5, and is calculated from *N_m_* using the following equation, with *k* denoting population size:

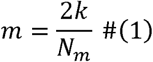

A total of 100,000 parameter sets are sampled. To filter for parameter sets with sufficiently beneficial resistance alleles, the selection coefficients of both resistance alleles (*S_A_,S_B_*) are estimated analytically by comparing the relative fitness of the heterozygote genotype (*ω_Aa_,ω_Bb_*) to the respective homozygote susceptible genotype (*ω_aa_,ω_bb_*):

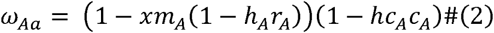

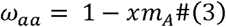

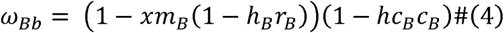

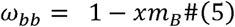

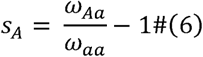

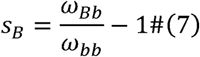

With estimated selection coefficients, the number of generations required for resistance to reach 50% in the population assuming it started at 1 copy prior to selection is calculated using continuous time approximation (Madgwick & Kanitz, 2022a). The first 10,000 parameters sets where both resistance alleles take less than 200 generations to reach 50% frequency by this estimate form the basis for all subsequent simulations.

### Simulating *de novo* mutation

distinct mutation transition matrices are constructed for diploid and haploid genotypes to simulate *de novo* mutations. The construction of the mutation transition matrices assumes that each haploid allele copy has a probability *m* to mutate, and a probability 1-*m* not to mutate. The probability of transitioning between any two genotypes is then calculated based on the number of alleles required to mutate and the number of ways to achieve the desired genotype (e.g. the probability of transitioning from the fully susceptible *aabb* genotype to the fully resistant *AABB* genotype is *m*^4^, as all four alleles must mutate and this is the only way to achieve such a transition). Each haploid allele copy can mutate at most once per generation.

### Simulating distinct reproductive strategies

four models are implemented to represent different reproductive strategies consistent with discrete-generation Wright-Fisher dynamics. Genotype frequencies are explicitly tracked, and selection is imposed prior to mating.

- The **sexual reproduction** (SR) model forms diploid zygotes from parental gametes, and *de novo* mutation is simulated using the diploid mutation transition matrix on zygote frequencies. Multinomial sampling on the post-mutation genotype distribution gave rise to the offspring genotype distribution, incorporating stochasticity. The SR model acts as a benchmark for assessing the influence of reproductive mode on resistance evolution.
- The **obligate parthenogenesis** or asexual reproduction (ASR) model forms diploid offspring by cloning parental genotypes. Diploid *de novo* mutation and multinomial sampling are applied to the normalised offspring genotype distribution. ASR is comparatively rare in insect pests, being observed in some localised aphid populations (Harrison & Mondor, 2011; Thia et al., 2025).
- The **cyclical parthenogenesis** (CYC) model is a combination SR and ASR dynamics. As insects employing cyclical parthenogenesis typically mate once per season (Dedryver et al., 2013), the CYC model follows SR dynamics for the last generation of each cycle. ASR dynamics occur on all other generations. CYC is common in aphid species (Margaritopoulos et al., 2009). To cover a range of aphid reproductive rates, two cycle lengths are simulated, lasting either 10 or 25 generations.
- The **haplodiploid** (HPD) model tracks 13 genotypes: nine diploid female and four haploid male. Selection and gamete formation are performed separately for each sex, with gamete frequencies normalised within sexes. Female zygotes are formed from both parental gametes, whereas male offspring arise solely from maternal gametes. Diploid mutation is applied to female zygotes and haploid mutation to male offspring, after which multinomial sampling generates offspring populations with the predetermined sex ratio. Examples of HPD pests include Western flower thrips (Frankliniella occidentalis) and silverleaf whiteflies (*Bemisia tabaci*). Sex ratio in Western flower thrips is biased towards females (Ding et al., 2018) and closer to parity in silverleaf whiteflies (Vyskočilová et al., 2019). To cover this range, two sex ratios are simulated, with females comprising either 50% or 75% of the population.

For each reproductive mode, separate simulations are run for sequences and mixtures. Sequences begin with compound A then upon its failure switch to compound B. Mixtures employ both compounds simultaneously throughout the simulation.

### Simulation termination and output

the simulation terminates if both resistance allele frequencies have reached the failure threshold, otherwise it proceeds until generation 2000 at which point the simulation terminates unconditionally. Within each simulation, three endpoint values are collected for each resistance allele:

- **Time to heterozygote emergence**: the number of generations lapsed until the stable establishment of the resistant heterozygote (i.e. no longer lost from the population).
- **Time to homozygote emergence**: the number of generations lapsed until the stable establishment of the resistant homozygote (i.e. no longer lost from the population).
- **Time to compound failure**: the number of generations lapsed until the population resistance allele frequency first reached the predetermined failure threshold. For sequences, compound A failure triggers switching to compound B. For mixtures, failure of one compound has no effect on the application regimen.

### Difference distribution analysis

comparisons across compound strategies (sequences as the reference against mixtures) and reproductive modes (sexual reproduction as the reference) are first made for each identical parameter set, then aggregated for all parameter sets. Within a parameter set, each comparison consists of randomly drawing one simulation per condition to be compared and calculating the difference (in generations) of the outcome of interest. This process is repeated 10,000 times per parameter set to account for stochastic variation among simulation replicates. Collecting results across all parameter sets generates the overall difference distribution.

## Results

### Mixtures are not disfavoured in asexually reproducing populations

The most direct measure of the effectiveness of a resistance management strategy is the time taken for a compound or a strategy to fail. First, to assess the overall dynamics across reproductive modes, the number of generations taken for compound A (the first compound applied in sequences) to fail are compared, with sexual reproduction (SR) model outcomes as reference. Using 25% as the failure threshold, when sequences are applied, no reproductive modes displayed a significant disadvantage compared to SR (Fig. 1a, Supp. 1a). There is a tendency for haplodiploid populations to reach the failure threshold faster than SR. All other reproductive modes show no observable difference in failure time distribution when compared to SR. Apart from haplodiploid populations, resistance evolution dynamics under sequences application does not seem to be strongly influenced by reproductive mode.

**Fig 1.**
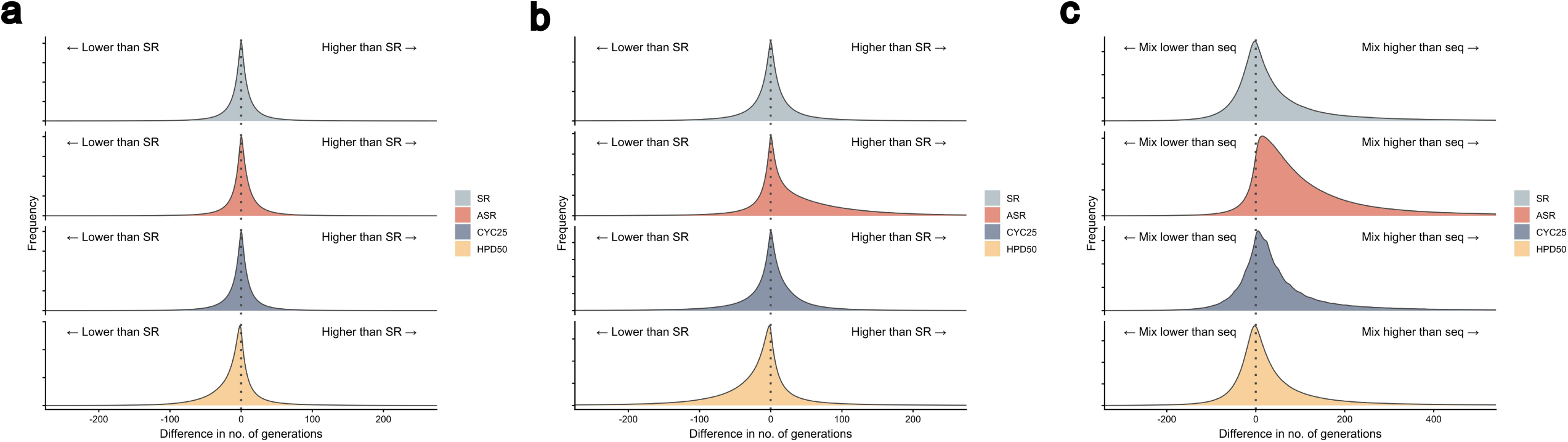
Comparison of compound and strategy failure times across reproductive modes. 25% failure threshold. a) Sequences, cumulative frequency distributions (CFD) of the difference in number of generations taken for resistance allele A to reach failure threshold. b) Mixtures, CFD of the difference in number of generations taken for resistance allele A to reach failure threshold. c) CFD comparing strategy failure times within each reproductive mode, with sequences as reference. SR – sexual reproduction, ASR – obligate parthenogenesis or asexual reproduction, CYC25 – cyclical parthenogenesis with one mating event every 25 generations, HPD50 – haplodiploidy with 50% females.

When mixtures are applied, with 25% as the failure threshold, similar trends are observed. Compared to sequences, mixtures result in higher variances in simulation outcomes for all reproductive modes (Fig. 1b, Supp. 1b). Resistance evolution is faster in haplodiploid populations as with sequences, indicating a strategic-agnostic mechanism. In some populations which reproduce exclusively through parthenogenesis (ASR in figure), resistance evolution is slower than SR. This suggests that the concurrent application of two compounds as a mixture caused competition between the two monogenic resistance alleles as heterozygotes arising in different genetic backgrounds. The resulting clonal interference delayed resistance evolution. Curiously, despite infrequent mating in cyclical parthenogenetic populations, the same delay is not observed.

To directly compare the favourability of mixtures against sequences within each reproductive mode, the time taken for each strategy to fail in controlling both resistance alleles is compared. With a 25% failure threshold, haplodiploid populations produced cumulative frequency distributions which are the most similar to the reference distribution produced by SR (Fig. 1c, Supp. 1c). Cyclical parthenogenetic populations have a slight tendency to favour mixtures, whereas clonal populations strongly favour mixtures over sequences. This result corroborates the observation in Fig. 1b, where mixtures prevent resistance spread by causing clonal interference, a phenomenon not possible when compounds are applied sequentially. Overall, mixtures do not seem to be disfavoured in any simulated reproductive modes compared to SR, and may additionally be favourable for controlling resistance in ASR populations by causing clonal interference.

### Resistance evolution under clonal reproduction is severely restricted by homozygote formation

The presence of clonal interference between resistance alleles as heterozygotes in ASR populations prompted the question of whether the same process also delays homozygote formation. To assess the dynamics of homozygote formation, simulation outcomes at the 75% failure threshold are analysed. Reaching this threshold requires at least half the population to be homozygous resistant. In sequences where compound A is deployed continuously until failure, parthenogenesis delayed both homozygote emergence and its selection (Fig. 2a). The delayed emergence can be explained by the need for two *de novo* mutations to occur within the same lineage. However, the delay in homozygous selection cannot be similarly explained. It is likely that given the population is bottlenecked by homozygote formation, homozygous individuals, when they eventually emerge, must compete against a fully heterozygous resistant population. The selection coefficient for the homozygous resistant genotype is therefore reduced compared to a sexually reproducing population with genotypes distributions near Hardy-Weinberg equilibrium.

**Fig 2.**
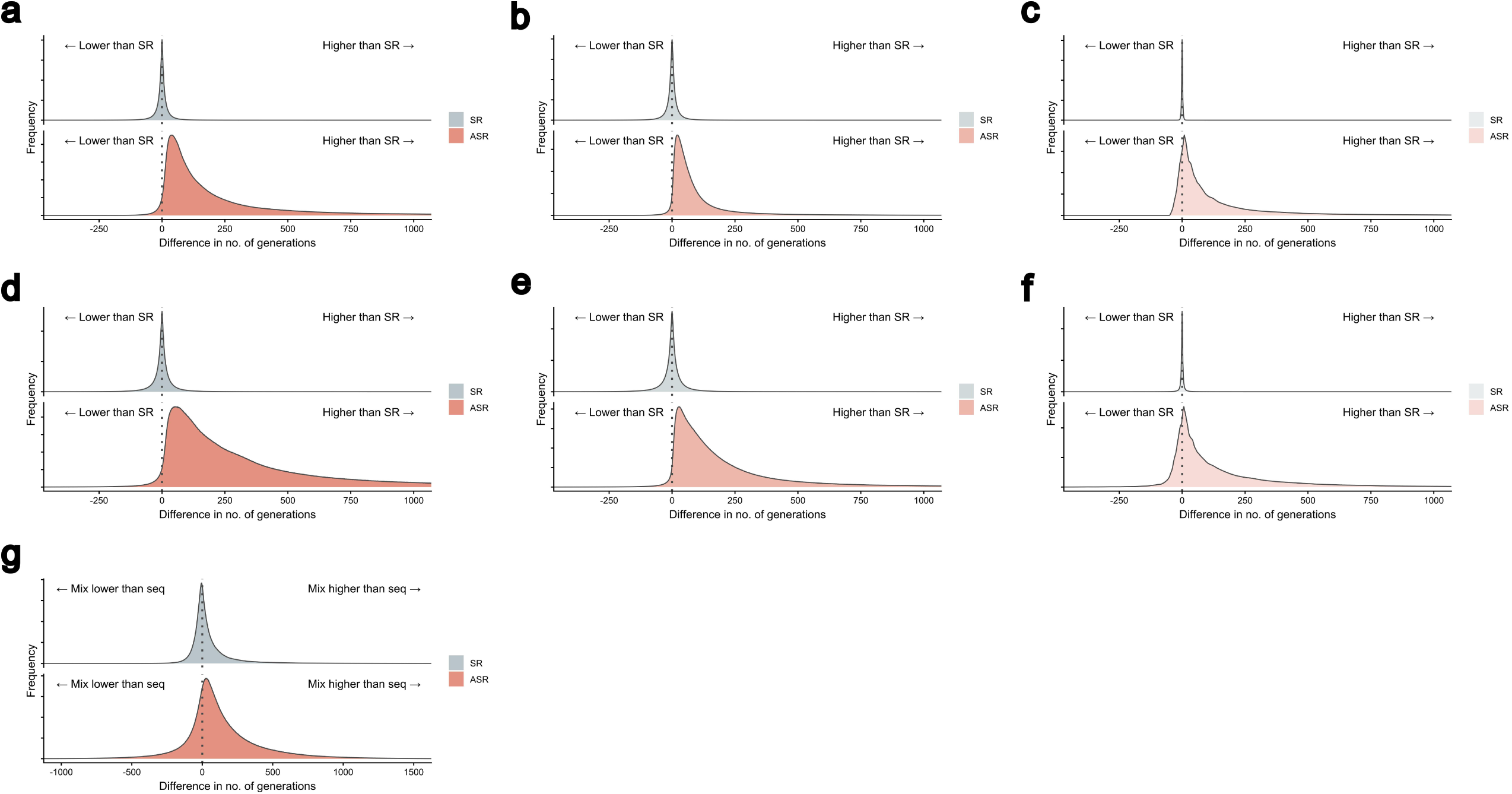
Homozygous resistance dynamics in obligate parthenogenesis. 75% failure threshold. a) Sequences, cumulative frequency distributions (CFD) of the difference in total time to compound failure. b) Sequences, CFD of the difference in homozygote emergence phase length. c) Sequences, CFD of the difference in homozygote selection phase length. d) Mixtures, CFD of the difference in total time to compound failure. e) Mixtures, CFD of the difference in homozygote emergence phase length. f) Mixtures, CFD of the difference in homozygote selection phase length. g) CFD comparing strategy failure times within each reproductive mode, with sequences as reference. SR – sexual reproduction, ASR – obligate parthenogenesis or asexual reproduction.

The same effect, whereby homozygote formation bottlenecks resistance evolution, is also observed when applying mixtures, and is generally more prominent compared to sequences (Fig. 2b). Mixtures further delay homozygote formation and spread due to clonal interference between the resistant homozygotes in addition to the resistant heterozygotes. The formation of the fully double-resistant individual requires four *de novo* mutation events to occur in the same lineage. The magnitude of this delay in adaptation is even more significant in the context of the parameter space, where only sufficiently beneficial alleles able to reach 50% frequency within 200 generations are chosen for simulation. Altogether, these effects result in mixtures being a much more favourable resistance management strategy at a 75% failure threshold than sequences (Fig. 2c).

### Rare mating events in cyclical parthenogenesis allow populations to rapidly adapt

With the exception of some localised aphid populations (Harrison & Mondor, 2011; Thia et al., 2025), few insect pests reproduce entirely through parthenogenesis. Cyclical parthenogenesis is much more common (Margaritopoulos et al., 2009). To probe the effect of rare mating events on resistance evolution in cyclical parthenogenesis, homozygote emergence times between cyclical parthenogenetic and sexually reproducing populations are compared. One mating event in every 10 generations is almost sufficient to completely remove the barrier introduced by homozygote formation in clonal reproduction when applying sequences (Fig. 3a) or mixtures (Fig. 3b). A small delay can be observed when one mating event occurs every 25 generations, but the magnitude of this delay is not nearly as large as in ASR populations. Across strategies, mixtures are able to delay the emergence of homozygous resistant individuals (Fig. 3c, 3d), but these individuals by large only emerge at regular intervals as dictated by the reproductive scheme, presumably on the rare mating generations. Overall, simulations indicate that rare mating events in cyclical parthenogenesis are able to effectively rescue the population from clonal interference, and allow more rapid resistance evolution with dynamics more closely resembling sexually reproducing populations.

**Fig 3.**
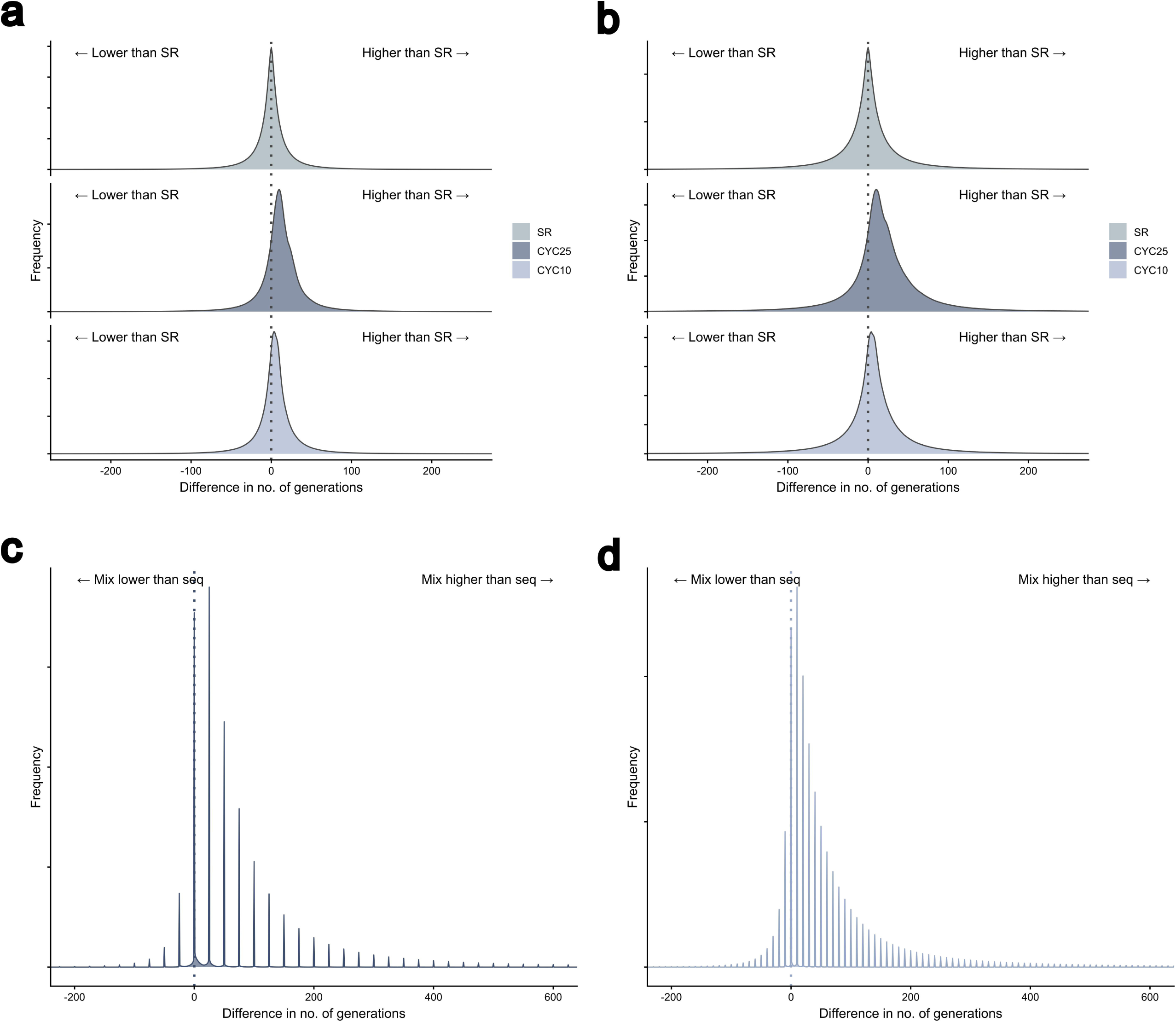
Homozygous resistance dynamics in cyclical parthenogenesis. 75% failure threshold. a) Sequences, cumulative frequency distributions (CFD) of the difference in number of generations taken for resistant homozygotes to emerge against compound A. b) Mixtures, CFD of the difference in number of generations taken for resistant homozygotes to emerge against compound A. c) CYC25, strategy comparison CFD of the difference in number of generations taken for resistant homozygotes to emerge, with sequences as reference. d) CYC10, strategy comparison CFD of the difference in number of generations taken for resistant homozygotes to emerge, with sequences as reference. SR – sexual reproduction, CYC25 – cyclical parthenogenesis with one mating event every 25 generations, CYC10 – cyclical parthenogenesis with one mating event every 10 generations.

### Resistance allele selection is faster in haplodiploid populations

In Fig. 1, haplodiploid populations have demonstrated a tendency to accelerate compound failure. This observation deserves more analysis to determine whether it results from a shortened emergence or selection phase. For both strategies, emergence of the heterozygote resistant genotype takes about the same length of time (or slightly longer) as an otherwise equivalent sexually reproducing population (Fig. 4a, 4b, Supp. 2a, 2b, top panels). However, the length of the heterozygote selection phase is almost always shorter in haplodiploid populations (Fig. 4a, 4b, Supp. 2a, 2b, bottom panels). Both effects are more pronounced in mixtures than sequences. Homozygous emergence and selection display a similar pattern (Supp. 3). Assuming the worst-case scenario where haploid resistant males are as fit as diploid resistant females, mixtures may be disfavoured on grounds of speeding up resistance allele selection in haplodiploid populations more so than sequences. Nonetheless, this effect did not clearly alter the aggregate favourability of mixtures in Fig 1c, which showed that haplodiploid populations have similar strategy preferences overall compared to sexually mating populations.

**Fig 4.**
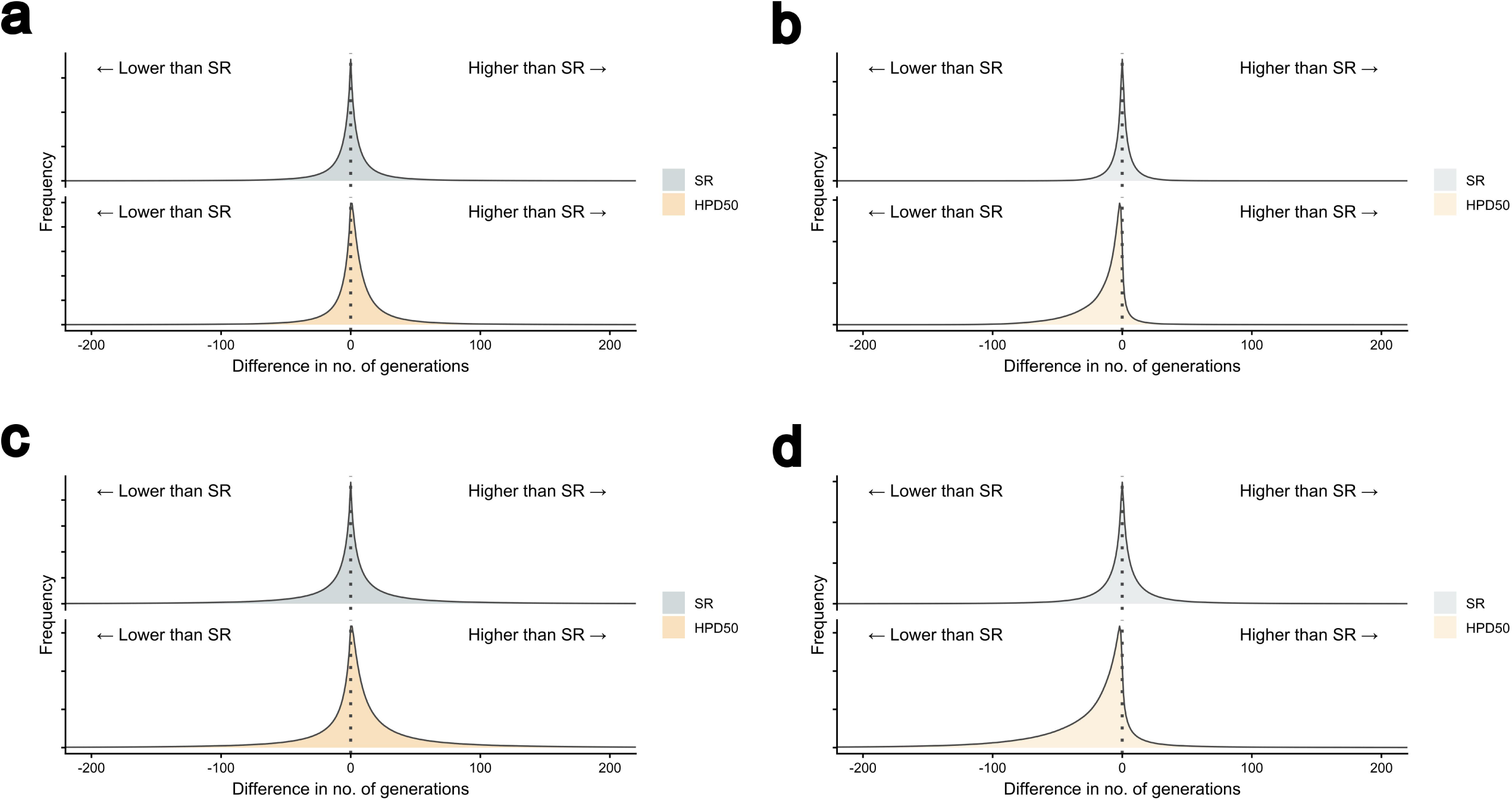
Heterozygote emergence and selection phase dynamics in haplodiploidy. 25% failure threshold. a) Sequences, cumulative frequency distribution (CFD) of the difference in heterozygote emergence phase length. b) Sequences, CFD of the difference in heterozygote selection phase length. c) Mixtures, CFD of the difference in heterozygote emergence phase length. d) Mixtures, CFD of the difference in heterozygote selection phase length. SR – sexual reproduction, HPD50 – haplodiploidy with 50% females.

## Discussion

Through extensive computational simulations, this work demonstrates that mixtures are generally not disadvantaged as a resistance management strategy over sequences on the basis of reproductive strategies. Among all asexual reproductive modes simulated, mixtures are by large at least as effective as in sexually reproducing populations. Mixtures are especially favourable in obligate parthenogenetic populations for its ability to induce clonal interference (between resistance alleles, as well as within each resistance allele in preventing homozygote formation) and delay resistance evolution. This clonal interference effect is much diminished in populations which reproduce through cyclical parthenogenesis, where rare mating events permit genetic recombination. Haplodiploid populations exhibit dynamics which generally resemble sexual reproduction, albeit with accelerated spread of resistance alleles due to haploid males, in agreement with prior theoretical literature (Crow & Kimura, 1965; Hedrick & Parker, 1997). With broadly similar dynamics observed across all reproductive modes, parameter space favourable for mixture deployment in mating populations, as determined by prior literature (REX Consortium, 2013; Levick et al., 2017; Madgwick & Kanitz, 2022b), is not expected to be significantly different in asexual pests.

These findings contrast with previous resistance modelling work which focused on selection phase dynamics (Helps et al., 2020). By excluding the emergence phase, such models implicitly assume that sexual and clonal populations enter selection with comparable genotype distributions. However, as demonstrated here, the absence of genetic recombination in clonal populations prevents the formation of certain genotypes, especially those carrying multiple copies of resistance alleles. Clonal populations therefore should have access to fewer genotypes than an equivalent mating population (Felsenstein, 1974), and assuming otherwise leads to biased predictions about how resistance will evolve. Accounting for emergence phase dynamics resolves this bias and leads to markedly different conclusions on long-term resistance evolution outcomes.

The present findings depend on evolutionary simulations which does not explicitly address companion traits frequently associated with asexuality. Parthenogenesis trades off the lack of genetic recombination with the ability to rapidly colonise and populate an environmental niche (Fujita et al., 2020), as these reproductive strategies do not depend on the costly process of finding a mate (Daly, 1978). This gives parthenogenetic populations a short-term demographic advantage, allowing larger populations to be attained quickly. Assuming a constant mutation rate, a larger population increases mutation supply, and this may offset some of the benefits of resistance management strategies by shortening the allele emergence phase. The demographic advantages of parthenogenesis may also alter the evolutionary timescale by shortening the length of each generation. The peach potato aphid is able to produce over 20 generations per year under favourable climates (van Emden et al., 1969; Forchibe et al., 2023), whereas sexually mating insect pests such as the codling moth, *Cydia pomonella*, may only have one to three generations within the same timeframe (Stoeckli et al., 2012). Nevertheless, differences in generation time should affect sequences as much as mixtures, hence this factor alone should not impact the overall strategy preference. One ecological factor that may differentially influence resistance management is the life-history traits of certain pest species. In pests such as thrips and aphids, limited dispersal due to the absence of winged individuals alters compound exposure patterns (Sudo et al., 2017). Resistant lineages may therefore remain clustered in treated areas, increasing the chance of acquiring additional resistance mutations under continued selection (Liu & Weissman, 2026).

Broadly, these results do not support general claims that asexual pests require different strategies for resistance management than sexual pests, resolving the discrepancy with empirical and theoretical evidence from other clonal systems (Arts & Hazuda, 2012; Bayat Mokhtari et al., 2017; Tyers & Wright, 2019; Jin et al., 2023). In other clonal systems, mixtures are able to delay resistance evolution by restricting the emergence multi-resistant clones (MacLean et al., 2010; McGranahan & Swanton, 2017; Labrie et al., 2022). As shown in this work, the same principle applies to insects which reproduce through obligate parthenogenesis. The presence of genetic recombination, even at low frequencies, greatly diminishes this effect and restores resistance evolution dynamics similar to what is seen in sexually reproducing populations. With the exception of obligate parthenogenetic pests, where mixtures have a unique advantage, the dynamics of resistance evolution in pests with alternative reproductive modes resemble their obligately mating counterparts.

## Data availability statement

All simulation code and data can be found on Zenodo: https://zenodo.org/records/22043568.

## Acknowledgement

This work is supported by funding from the Biotechnology and Biological Sciences Research Council, grant number BB/T008784/1, a Magdalen Graduate Scholarship in Biology, and Syngenta Limited.

