## Supplementary material for "Evaluating pesticide mixtures for resistance management in asexual insect pests": Table 1

Table 1. Simulation parameters and respective ranges.

| **Parameter** | **Symbol** | **Range** |
| --- | --- | --- |
| Population size | $k$ | 10^4^ – 10^9^ |
| Compound exposure | $x$ | 0 – 1 |
| Compound efficacy | $m_{A}, m_{B}$ | 0 – 1 |
| Recovery conferred by resistance allele | $r_{A}, r_{B}$ | 0 – 1 |
| Dominance of resistance allele | $h_{A}, h_{B}$ | 0 – 1 |
| Cost of resistance allele | $c_{A}, c_{B}$ | 10^-3^ – 10^-0.5^ |
| Dominance of cost of resistance allele | $hc_{A}, hc_{B}$ | 0 – 1 |
