## Supplementary figures and legends for "Evaluating pesticide mixtures for resistance management in asexual insect pests"

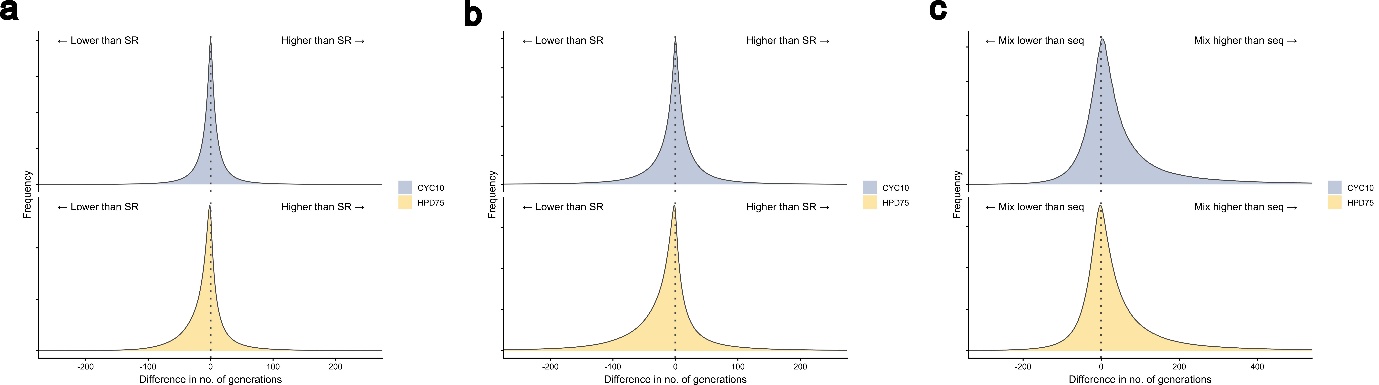

**Supp 1.** **Comparison of compound and strategy failure times across alternative reproductive modes.** 25% failure threshold. a) Sequences, cumulative frequency distribution (CFD) of the difference in number of generations taken for resistance allele A to reach failure threshold. b) Mixtures, CFD of the difference in number of generations taken for resistance allele A to reach failure threshold. c) CFD comparing strategy failure times within each reproductive mode, with sequences as reference. CYC10 – cyclical parthenogenesis with one mating event every 10 generations, HPD75 – haplodiploidy with 75% females.

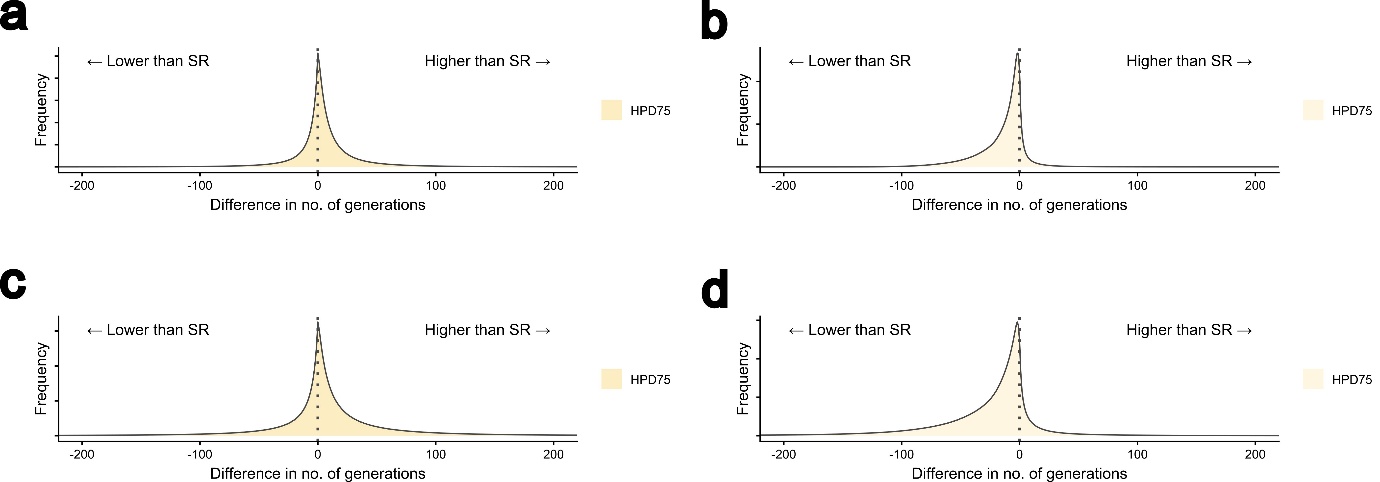

**Supp 2. Heterozygote emergence and selection phase dynamics in HPD75.** Cumulative frequency distributions of the difference in heterozygote emergence phase length and heterozygote selection phase length, with 25% as failure threshold. a) Sequences, heterozygote emergence phase length. b) Sequences, heterozygote selection phase length. c) Mixtures, heterozygote emergence phase length. d) Mixtures, heterozygote selection phase length. HPD75 – haplodiploidy with 75% females.

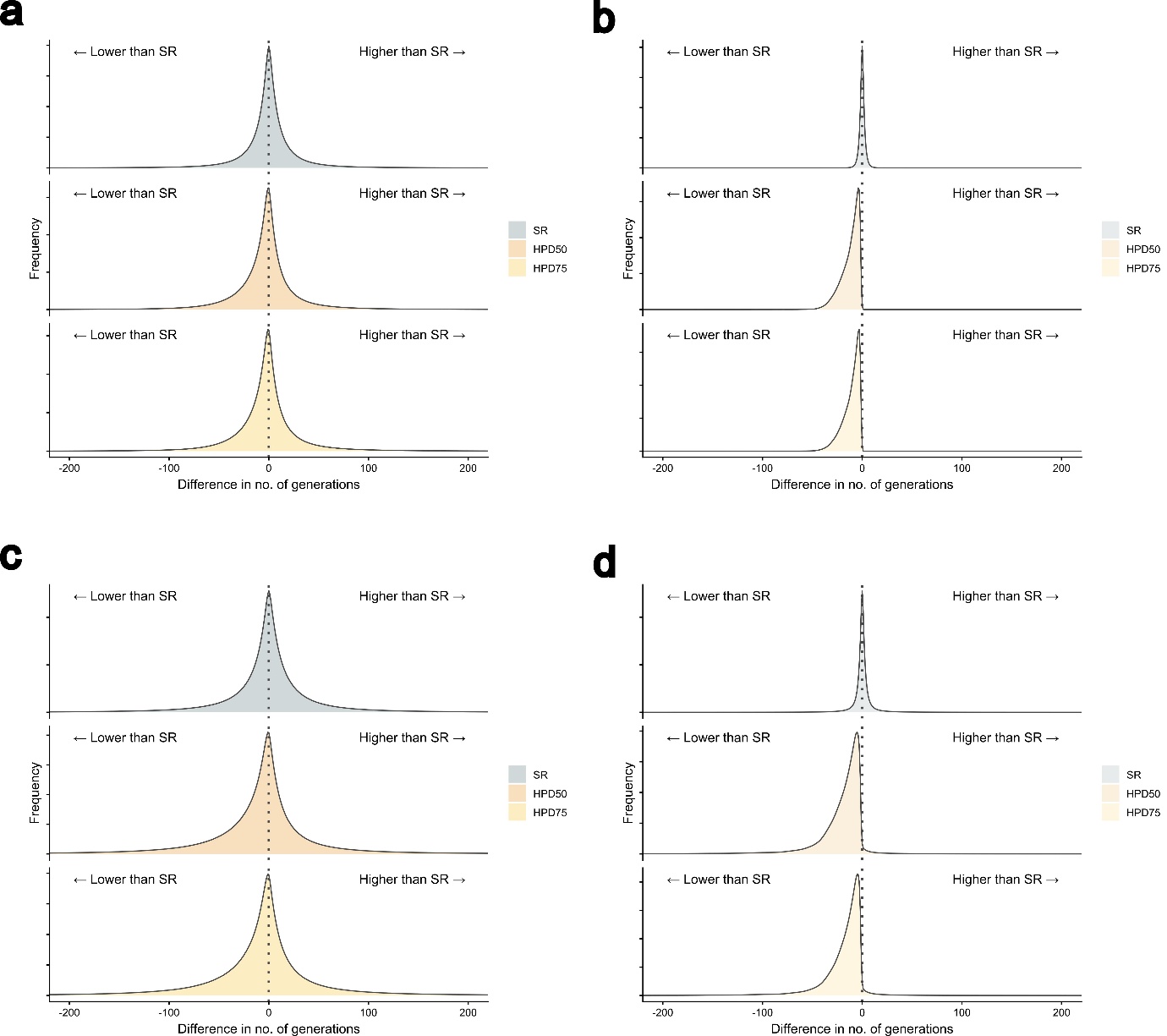

**Supp 3. Homozygote emergence and selection phase dynamics in haplodiploidy.** Cumulative frequency distributions of the difference in homozygote emergence phase length and homozygote selection phase length, with 75% as failure threshold. a) Sequences, homozygote emergence phase length. b) Sequences, homozygote selection phase length. c) Mixtures, homozygote emergence phase length. d) Mixtures, homozygote selection phase length. SR – sexual reproduction, HPD50 – haplodiploidy with 50% females. HPD75 – haplodiploidy with 75% females.
